# EZ-SPOTs: A simple and robust high-throughput liquid handling platform

**DOI:** 10.1101/2024.05.13.594031

**Authors:** Jon Albo, Shenghao Tan, Joee D Denis, Rebecca Franklin-Guild, Samira Shiri, Kelsi M Sandoz, Nate J Cira

**Affiliations:** Meinig School of Biomedical Engineering, Cornell University, Ithaca, New York, 14853, United States; Department of Microbiology and Immunology, Cornell University, Ithaca, New York, 14853, United States; Department of Population Medicine and Diagnostic Medicine, Animal Health Diagnostic Center, College of Veterinary Medicine, Cornell University, Ithaca, New York, 14853, United States; Department of Mechanical Engineering, University of Utah, Salt Lake City, Utah, 84112, United States

## Abstract

Liquid handling is a fundamental capability for many scientific experiments. Previously, we introduced the Surface Patterned Omniphobic Tiles (SPOTs) platform, which enables manipulation of hundreds to thousands of independent experiments without costly equipment or excessive consumable expenses. However, the SPOTs platform requires a custom coating formulation and lacks robustness. To overcome these limitations, we introduce EZ-SPOTs. These devices can be created in an hour with common fabrication tools and just three components - glass, a hydrophobic coating, and acrylic. EZ-SPOTs preserve many of the SPOTs platform’s strengths - ease of use, ability to handle a wide range of volumes, and scalability - and adopt a durable and abrasion resistant coating that enables multiple reuses of each device. Here, we describe the fabrication of EZ-SPOTs and showcase how its reusability allows antibiotic susceptibility testing of many isolates using a single device. These results quantitatively match current gold standard assays and the increased throughput provides substantially more information than standard approaches.

## 1. Introduction

Liquid handling is a routine task critical to laboratory procedures in life sciences, healthcare, and medical research. Traditional micropipetting is a versatile method for handling liquids in a broad range of volumes but is limited in scalability, requires large sample volumes, has a high risk of human error, and cannot be automated for complex workflows. Pipetting robots address limitations of automation and human error but require high initial and maintenance costs and expertise to set-up and use. A wide array of microfluidic techniques have their own advantages, however, they are often application-limited or require specialized expertise and equipment. Emulsion-based droplet microfluidic techniques^1,2^, are highly efficient at rapidly generating and interrogating many thousands of individual reactions. However, they rely on two immiscible liquids, and modifying or extracting the droplets for downstream processes proves challenging. Droplet microarrays^3,4^, offers multiplexing^5,6^ and sample recovery^4^, but are restricted in compatible liquid types and lack precise loading mechanisms for discontinuous dewetting based systems.

Previously, we introduced the Surface Patterned Omniphobic Tiles (SPOTs) platform^7^, which builds on droplet microarrays and uses a superomniphobic coating^8,9^ that can repel liquids with low surface tensions. This platform allows for precise deposition of sub-microliter volumes of liquids using a passive loading device without the need for specialized fabrication equipment. We demonstrated its versatility and ease of use through diverse applications such as culturing bacterial cells, growing perovskite crystals, and reagent metering and addition for polymerase chain reaction (PCR). Despite its advantages, the custom-synthesized superomniphobic coating on the SPOTs platform is challenging to apply, subject to batch variation, and requires high-temperature baking for full effectiveness. The coating lacks robustness as it rubs off very easily, making repeated use, shipping, and extensive manipulation difficult.

In this paper, we introduce EZ Surface Patterned Omniphobic Tiles (EZ-SPOTs) which utilizes a commercially available coating to develop a robust, reusable, and easy-to-make solution for performing hundreds to thousands of simultaneous experiments. The abrasion resistance and ease of fabrication makes this variation more widely accessible to laboratories across the world. We then demonstrate the utility of the EZ-SPOTs platform by performing antibiotic susceptibility testing (ASTs) as a proof-of-concept.

## 2. Results and Discussion

### 2.1 The EZ-SPOTs Platform

The EZ-SPOTs platform contains two main components: 1) EZ-SPOTs plates that selectively repel and attract liquids in defined patterns (Fig. 1A), and 2) loading devices that accurately deposit liquids by sliding across EZ-SPOTs plates (Fig. 1B). The platform can be used by either designing the plates to interface directly with existing microtiter plates or performing experiments directly on the EZ-SPOTs plates where liquids are combined by sandwiching multiple plates together^5,6^.

**Figure 1.**
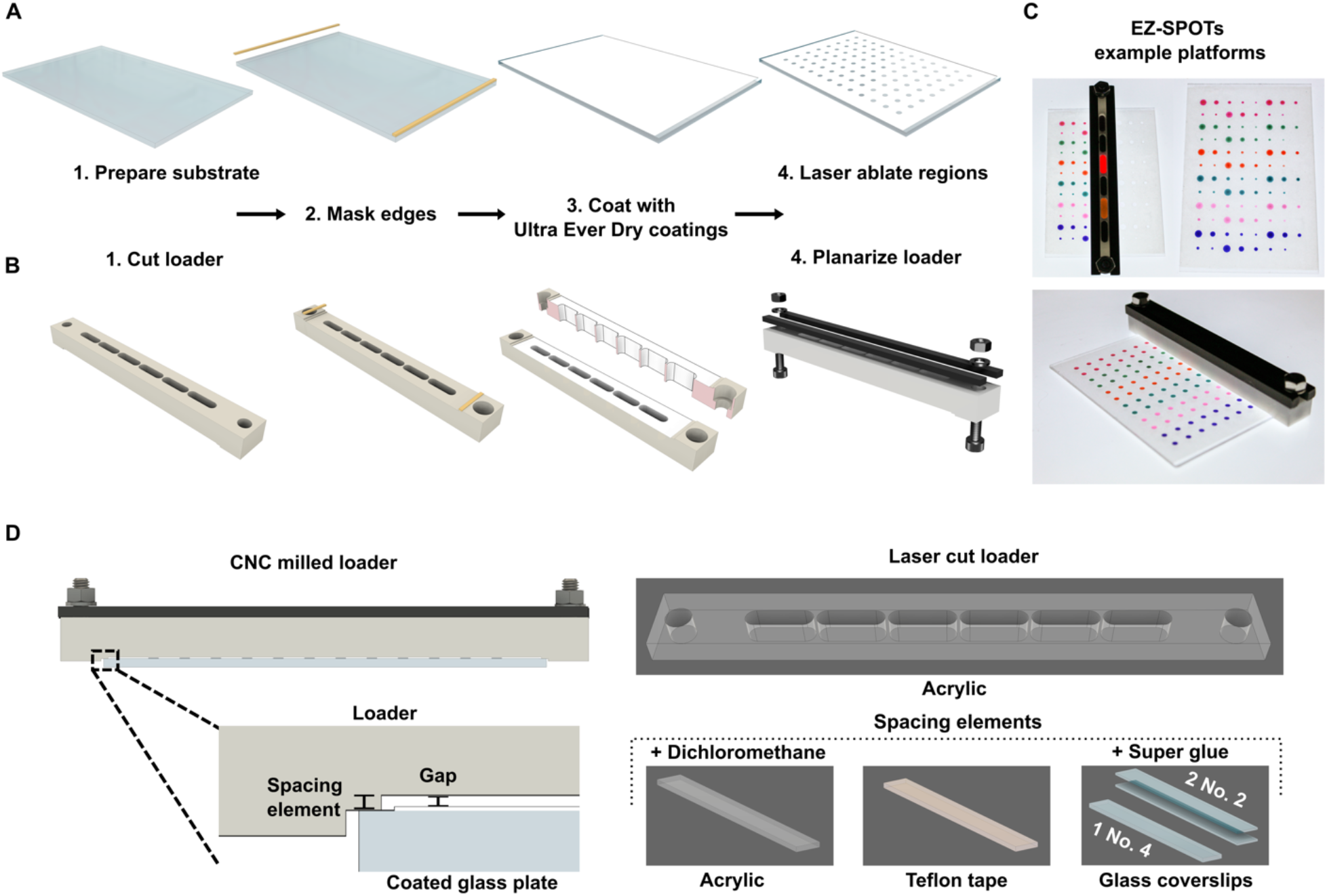
Fabrication of EZ-SPOTs platform. A) EZ-SPOTs plates are fabricated by cleaning the glass, masking the edges with tape to ensure smooth loader sliding, coat with Ultra Ever Dry bottom and top coatings, and laser ablate specific regions; B) EZ-SPOTs loaders are fabricated by cutting a loader, masking the edges with tape to ensure smooth sliding on the glass, coat with Ultra Ever Dry bottom and top coatings on the bottom and inside of the loader, and planarize the plastic to alleviate any deformities; C) Examples of different laser ablation patterns on EZ-SPOTs platforms; D) Schematics showing the important spacing characteristics of the loader indicating that the spacing element can be directly milled when using a CNC or can be adhered through different means if using a laser cutter.

To ensure that the EZ-SPOTs platform could be used in a wide range of experiments, we carefully selected a commercially available coating that met four key criteria: easy to apply, biocompatible, durable, and not requiring any custom synthesis. We tested many custom and commercially available coatings and chose Ultra-Ever Dry (UED) bottom and top coatings (UltraTech International) which fit all criteria and result in reusable plates and loaders. The platform can be fabricated anywhere with adequate airflow and a laser cutter as it does not require the use of lithography, synthesis of custom molecules, specialized equipment, or high temperature heat treatment.

#### 2.1.1 Fabrication of EZ-SPOTs plates and loaders

EZ-SPOTs plates are fabricated with just a few materials and a laser cutter (Fig. 1A). The fabrication process involves cleaning the glass surface, masking the edges with tape for smooth sliding, applying UED bottom and top coatings, and laser ablating specific regions to create -philic circles for holding liquid. The diameter of the ablated -philic regions can be varied to adjust the volume of liquid deposited.

EZ-SPOTs loaders are also fabricated with just a few materials and CNC machining or a laser cutter (Fig. 1B). The fabrication process involves cutting a predesigned loader from plastic stock, masking the edges with tape for smooth sliding, applying UED bottom and top coatings, and optionally planarizing the loader with metal bars to alleviate any plastic deformation. Each loader has one or more reservoirs to deposit liquids (typically a slot, slots, or holes) and spacers to create consistent gaps with the plate (Fig. 1D). In Fig. 1B, the loader has six slots to load the same liquid in every two columns of the 96 regions EZ-SPOTs plate (Fig. 1C). The size of the reservoir can be varied to fill different rows, columns, or sections of the plate. Loaders can also be fabricated using various materials and methods, including laser cutting and adhering spacers of consistent thicknesses (Fig. 1E).

### 2.2 Performance of EZ-SPOTs

We first evaluated the consistency and precision of EZ-SPOTs, SPOTs, and pipettes for handling volumes over two to three orders of magnitude. Variation was measured using the coefficient of variation (CV), CV= ^*σ*^/_*μ*_ *100% where *σ* is the standard deviation and *μ* is the mean. For a single device (Fig. 2A), both EZ-SPOTs and SPOTs had much lower CVs than a pipette and were able to maintain this consistency at smaller volumes. For multiple devices (Fig. 2B), pipettes have extremely high CVs at small volumes while EZ-SPOTs and SPOTs maintained comparably low CVs. The low CV demonstrated by EZ-SPOTs is crucial for experiments that require high precision. With the design characteristics described in this paper, the EZ-SPOTs platform can achieve a wide volume range of 40 nL to 6 µL in the diameter range of 0.5 mm to 4.5 mm (Fig 2C).

**Figure 2.**
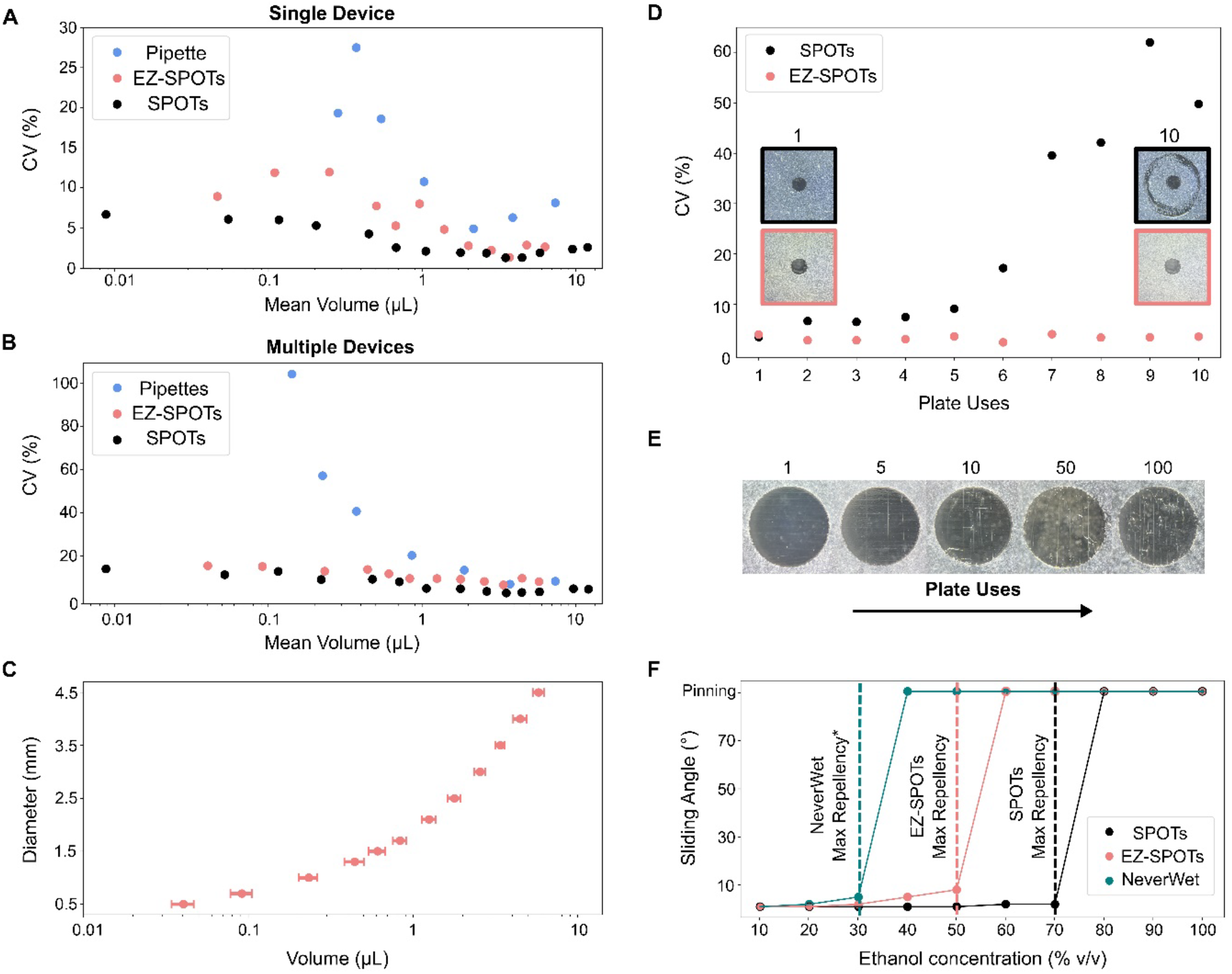
Performance of the EZ-SPOTs platform. A) Variation of deposited volumes (n=3) using a single pipette, a single EZ-SPOTs plate and loader, and a single SPOTs plate and loader; B) Variation of deposited volumes (n=3) using three pipettes, three EZ-SPOTs plates and loaders, and three SPOTs plates and loaders; C) Volume range for different diameters of laser-ablated areas using a specific gap height and laser ablation parameters; D) Comparison of the volume variation (n=8) between the EZ-SPOTs platform and the SPOTs platform when reusing the same plate 10 times; E) Additional uses of EZ-SPOTs plates results in increased microcracks within a single laser-ablated region; F) Comparison of the ethanol concentration (% v/v) that can be repelled by NeverWet (a superhydrophohic coating), EZ-SPOTs, and SPOTs. *NeverWet fails if water is left on it for an extended period of time and does not repel most biological liquids or DMSO. SPOTs and pipette data in A and B adapted from Shiri et al.7

To test the durability of EZ-SPOTs, we mimicked repeated use by interfacing the plates with 96-well microtiter plates (consisting of the coating rubbing against the raised edges of each well on the 96-well plates) and measuring the CV of eight metering sites on the plates after each cycle (Fig. 2D). We found that with multiple uses, the CV for the EZ-SPOTs plates remained consistently under 4%, while the SPOTs plates rapidly increased to over 60%. This rapid increase in CV is due to the SPOTs coating being damaged during liquid transfer to a microtiter plate, which exposes unwanted -philic areas that retain liquid and lead to increased volume. The durability of EZ-SPOTs is seen with the consistently low CVs, however, we did find that the laser ablation required to remove the UED coating creates microcracks on the glass surface which can propagate as number of uses increase (Fig. 2E). Fortunately, the performance is unaffected as these cracks do not affect volume deposition and any cells that migrate into the cracks do not survive when plates are soaked in bleach.

Lastly, we assessed the solvent compatibility by measuring the sliding angle of different solutions of ethanol-water concentrations on the coated surface (Fig. 2F). We observed that compatible liquids with the platform roll off with a low sliding angle while unsuitable liquids pin or wet the coating in less than 10 seconds. The EZ-SPOTs platform can repel up to 50% ethanol, which is better than superhydrophobic coatings like NeverWet that can only repel up to 30% (Fig. 2F). Although this repellency is less than the SPOTs platform, the EZ-SPOTs platform can still effectively repel many lower surface tension liquids including DMSO, PCR reaction mix, protein solutions, and many organic compounds.

### 2.3 Use of EZ-SPOTs for Antibiotic Susceptibility Testing (AST)

We validated the EZ-SPOTs platform with a real-world example by performing ASTs as they require conducting numerous repetitive assays with precise antibiotic concentrations. The EZ-SPOTs platform offers greater flexibility in the selection of drugs and concentrations, making it useful to test multidrug-resistant (MDR) organisms, or when commercial platforms are insufficient. We conducted ASTs on 12 clinical methicillin-resistant *Staphylococcus pseudintermedius* (MRSP) isolates using only two EZ-SPOTs plates to meter and transfer drugs and isolates to 96-well microtiter plates (Fig. 3A, 3B, 3C). MRSP isolates are an emerging pathogen in humans and animals (primarily dogs)^10–12^, and their ability to transmit between species^10,13,14^ makes it crucial to monitor resistance trends for a wide range of drugs and concentrations, including those not typically included on commercial AST panels. We tested six antibiotics, five of which are on the commercially available Thermo Fisher Sensititre panel at an expanded concentration range. The assay provides a way to obtain growth curves for drugs at many concentrations for isolates (Fig. 4A).

**Figure 3.**
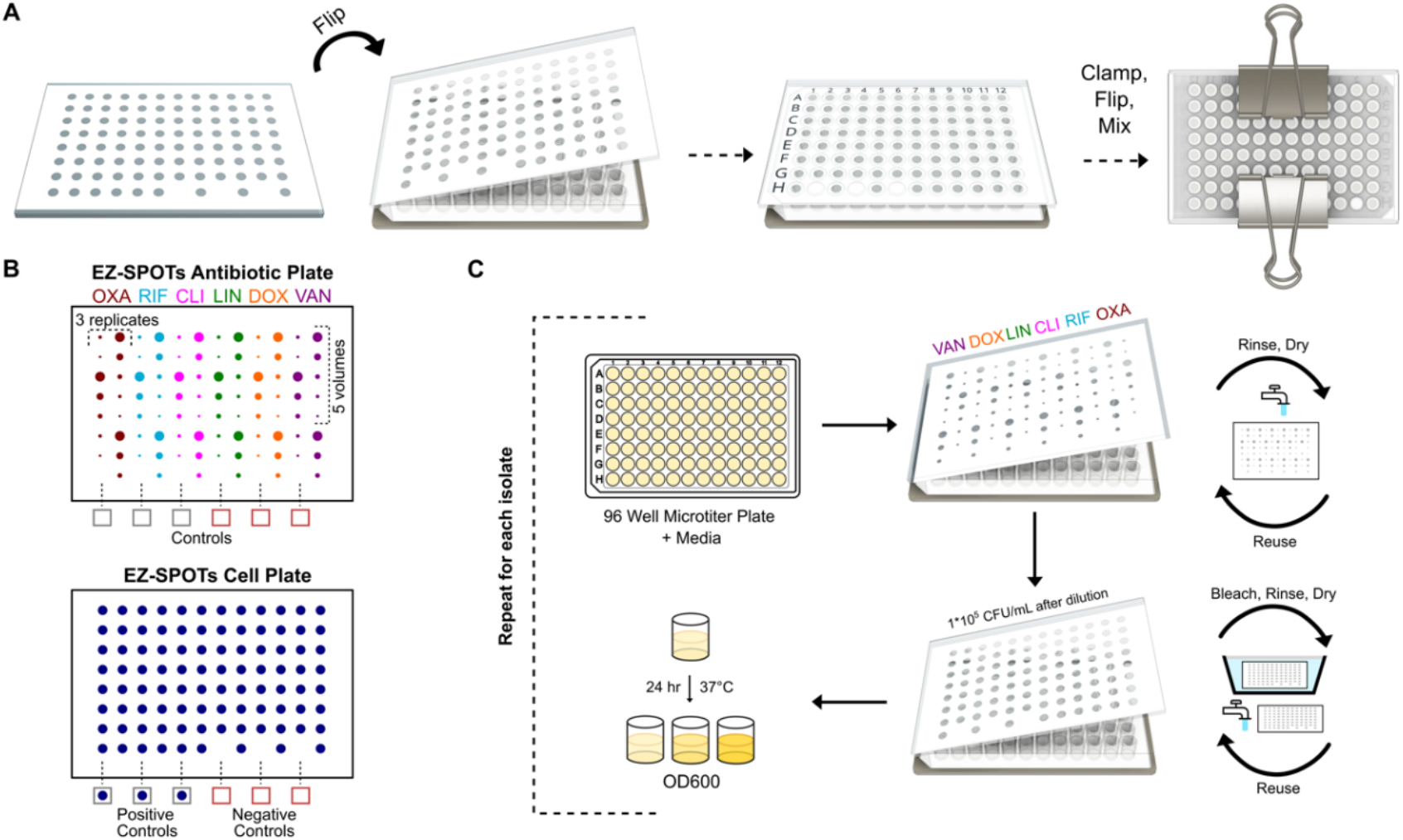
Experimental design using the EZ-SPOTs platform for ASTs. A) EZ-SPOTs plates are interfaced with 96-well microtiter plates by lining up and flipping the EZ-SPOTs plate on top of the 96-well microtiter plate, clamping two sides, flipping the two plates upside down, and shaking until well-mixed; B) Schematics of the EZ-SPOTs antibiotic plate design which contains three replicates of six antibiotics at five different volumes and six coated areas for controls and the EZ-SPOTs cell plate which is loaded with one isolate and has spaces for three positive controls and three negative controls; C) a 96-well microtiter plate was filled with media, interfaced with the EZ-SPOTs antibiotic plate which was loaded with a six-slot loader, then the resulting 96-well plate was interfaced with the EZ-SPOTs cell plate which was loaded with a single-slot loader and each 96-well microtiter plate was incubated until an OD600 reading. The EZ-SPOTs antibiotic plate was rinsed and reused for each isolate and the EZ-SPOTs cell plate was bleached for two minutes, rinsed, and reused for each isolate.

**Figure 4.**
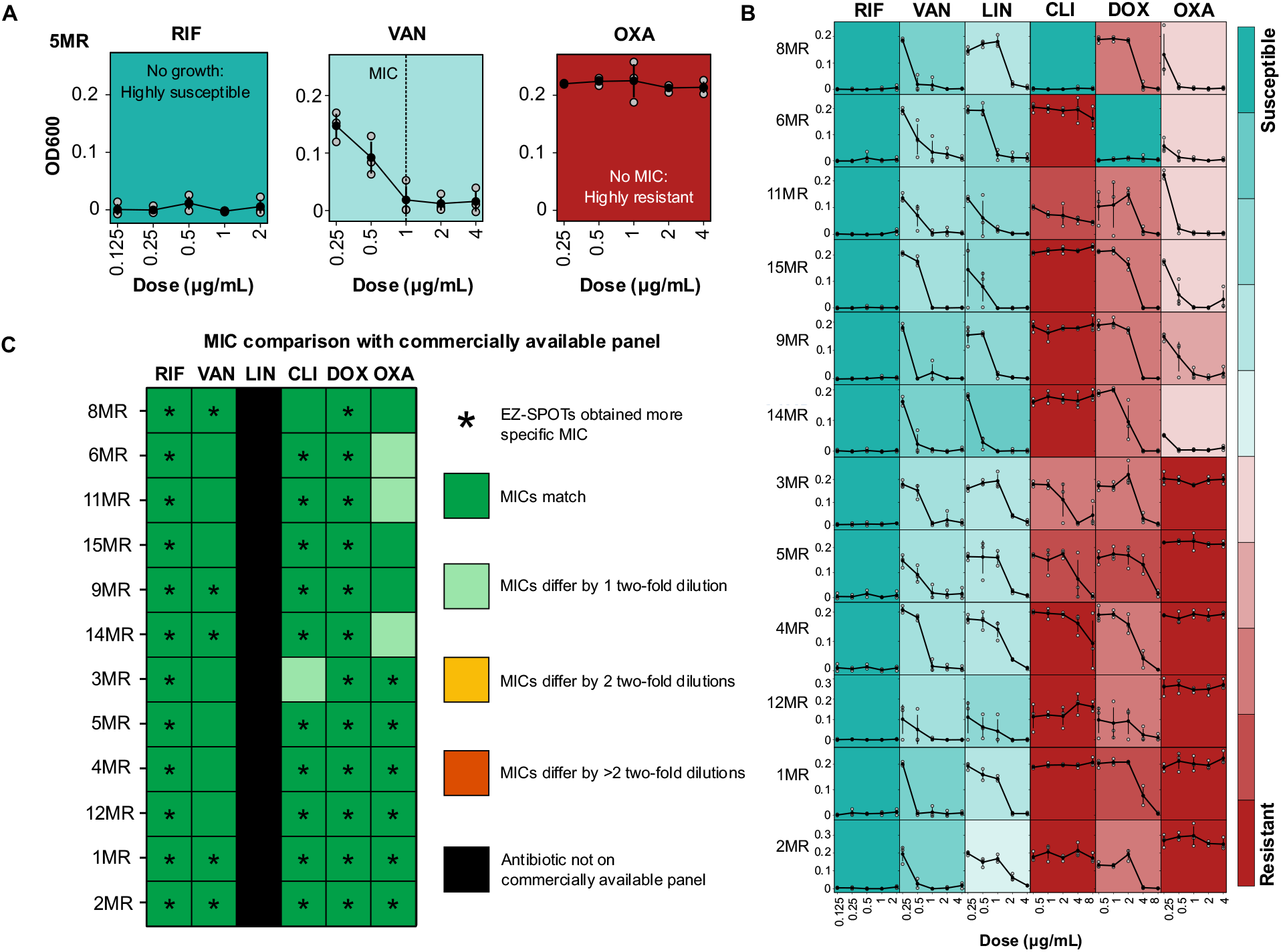
Results from AST experiments. A) Dose-response curves for a single isolate with a single antibiotic where gray points (n=3) indicate replicates for each dose and black points indicate averages with standard deviation bars. Background color is based on the MIC (defined as the lowest concentration of an antibiotic that completely inhibits growth) of each isolate and is scaled based on CLSI resistance or susceptibility breakpoints for each antibiotic; B) Dose-response curves for 12 isolates against 6 antibiotics where most are seen to be highly resistant to certain antibiotics and a range of susceptibilities to others; C) MIC comparison to commercially available Sensititre panel where most MIC’s match identically and few are only one two-fold dilution off. Additionally, more specific MIC information was obtained for most isolates and antibiotics.

Evaluating our assay revealed that 10 out of the 12 isolates were MDR as they were resistant to antibiotics from three or more classes (Fig. 4B). The isolates were found to be highly susceptible to rifampicin (RIF), susceptible to both vancomycin (VAN) and linezolid (LIN), mostly resistant to clindamycin (CLI) and doxycycline (DOX) apart from one isolate each, and all resistant to oxacillin (OXA). When comparing the MICs for each of the 72 isolate/antibiotic combinations with those obtained from the commercially available Sensititre panel (Fig. 4C), we found a near perfect agreement with only four instances of being one two-fold dilution different. Additionally, with the expanded concentration range, we were able to resolve more information about the true resistance and susceptibility of each isolate.

## 3. Materials and Methods

### Fabricating EZ-SPOTs plates

Borosilicate glass (127.76 mm x 85.48 mm) plates were first cleaned with acetone (VWR) and ethanol (VWR) to remove debris. Next, edges were masked (foam tape, 3.175 mm wide) based on the orientation of the loader to ensure a clean surface between the plate and the loader sliding on top. Then, Ultra-Ever Dry (UED) bottom coat was applied with four light coats, and while the bottom coat was still wet, the tape mask was removed. After at least half an hour, UED top coat (UltraTech International, batch B24) was applied with four light coats, the plates were left to dry at room temperature, and the plates were then rinsed with water to remove any loose coating. Current UED top coat batches may lack oleophobicity and only be hydrophobic instead of omniphobic. We recommend testing each reagent on the coating prior to use with a similar test to the sliding angle measurements. If an application requires more repellency than the UED top coating batch provides, the SPOTs coating^7^ can be applied as the top coating instead with similar results to UED top coat batch B24.

### Laser ablating coating to create -philic regions

Specific regions of the coating were laser ablated (Epilog Fusion Edge 40W, 1 cycle at 15% speed, 30% power followed by 1 cycle at 15% speed, 10% power) based on the experimental design. The laser settings must be consistent across a single experiment to ensure consistent volume deposition. Experiments are designed by using a spreadsheet software to plan desired volumes on each plate which are translated to diameters using volume characterization data. We then define the pitch between each diameter from the spreadsheet file in a MATLAB script and export the plates as an SVG file. Finally, we process this in Inkscape and send the file to the laser to selectively ablate the plates.

### Fabricating EZ-SPOTs loaders

When making a plastic (Delrin, McMaster Carr) loader, the loader was milled with desired slot, slots, or holes from the bottom side to create the spacer feature (0.4 mm thick) and ensure a consistent gap height. After milling, all milled areas were cleaned with a sharp blade. Next, the spacer feature was masked (foam tape, 3.175 mm wide) to ensure a clean surface between the loader and the plate.

When making a laser cut plastic (Acrylic, 6.35mm thick, McMaster Carr) loader, laser cut the loader with desired slot, slots, or holes. Spacer elements (0.254 mm acrylic, 0.5 mm Teflon tape, 1 No. 4 or 2 No. 2 glass coverslips) are cut and adhered (dichloromethane for acrylic or super glue for glass) to the laser cut plastic. Next, the spacer element is masked (foam tape, 3.175 mm wide) to ensure a clean surface between the loader and the plate.

Then for both the milled and laser cut loaders, UED bottom coat was applied with four light coats to all outer and inner surfaces of the loader, and while the bottom coat was still wet, the tape mask was removed to maintain a smooth surface. After at least half an hour, UED top coat (batch B24) was applied with six light coats to all outer and inner surfaces of the loader, and the loader was left to dry for at least a few hours or overnight. Finally, the loader was planarized (optional) with metal bars (precision-milled stainless steel, McMaster Carr), screws (Thor Labs), and nuts (Thor Labs) and tightened to ensure a consistent volume deposition across all areas of the plate. The loader was then slid over the EZ-SPOTs plate to remove top coating on the previously masked edges.

### Interfacing EZ-SPOTs plates with microtiter plates

EZ-SPOTs plates were interfaced with filled 96-well microtiter plates (VWR) by matching the 96-well plate layout. Then, the EZ-SPOTs plates were flipped over and lined up with the 96-well microtiter plates. Large binder clips were clamped on two sides, the combined plates were flipped over and shaken until well-mixed. The combined plates were flipped back over, given several shakes to ensure the liquid returned to the well in the 96-well microtiter plate, and the binder clips and EZ-SPOTs plate were removed. To move liquid from an EZ-SPOTs plate to an unloaded 96-well microtiter plates, the EZ-SPOTs plate would be matched to the 96-well plate, taped together, and put in a plate centrifuge at 700-2300 g for 30 seconds.

### Volume characterization

Volume deposition was dependent on both the gap height between the coated loader and the coated plate as well as the laser ablation parameters for the -philic regions on the EZ-SPOTs plate. To perform volume characterization, 96-well microtiter plates (VWR) were first loaded with 250 μL of DI water. A 24 g/L fluorescein solution was created and serial diluted by a micropipette into a 96-well microtiter plate for four replicates. Optical densities (OD) were obtained using a spectrophotometer (SpectraMax Plus 384 Microplate Reader, Molecular Devices) at OD470, OD520, and OD530. This was used to create linear graphs to relate the OD of fluorescein dilutions to volumes.

Next, the 24 g/L fluorescein solution was added to the loader and loaded into the -philic regions of an EZ-SPOTs plate with 12 diameters ranging from 0.5 mm to 4.5 mm and eight replicates of each diameter. This was performed on three EZ-SPOTs plates, repeated four times for each plate and we obtained OD470, OD520, and OD530 using a spectrophotometer. The OD readings were translated to volumes by using the equations from the linear graphs previously described. Each of these plates and replicates were used to create a calibration curve relating volume to diameter. This informed the diameter needed to be ablated to obtain a specific volume of liquid. This process was repeated for each different style of loader used.

### Volume variations of EZ-SPOTs, SPOTs, and pipettes

To determine the volume variation, we tested the loaded volume for a single device of each platform and for four EZ-SPOTs platforms, four SPOTs platforms, and three pipettes. The data displayed for SPOTs and pipettes was shown previously^7^, and here, we added the EZ-SPOTs volume variation. We performed this test by loading different volumes of a 24 g/L fluorescein solution into 96-well microtiter plates for each platform and then backing out the actual volume as previously described. We had 12 replicates for each pipette run, and 8 replicates for each EZ-SPOTs and SPOTs runs.

### Reusing EZ-SPOTs and SPOTs plates

To mimic repeated use of both the EZ-SPOTs and SPOTs plates, we followed our volume characterization procedure but repeated the characterization with the same plates 10 times. The plates were rinsed and the SPOTs plate was dried with compressed air while the EZ-SPOTs plate was dried by blotting it on a paper towel. Eight replicates of 3 mm diameter -philic regions were averaged for each point.

### Sliding angle measurements

The sliding angle was measured by dropping 10 μL of different ethanol-water concentrations 5 cm above a coated glass slide. If the liquid did not roll off the glass, the angle was increased one degree and the liquid was dropped again. The angle was increased until the liquid pinned to the surface or at the point at which the liquid could not be on the surface for more than 5 seconds without wetting the coating.

### Bacterial calibration curve

To relate an OD600 reading to bacterial cell concentration, three MRSP isolates (Cornell Animal Health Diagnostic Center, Bacteriology Laboratory) were grown with and without shaking at 37°C in cation-adjusted Mueller Hinton broth (CAMHB) for 24 hours. A spectrophotometer was used to obtain the OD600 for each suspension. Then, serial dilutions were made for each suspension ranging from 10^-3^ to 10^-7^ and 100 μL of each of these were streaked on a 5% sheep’s blood in tryptic soy agar plate (VWR) and incubated at 37°C for 24 hours. Finally, plates containing between 30 and 300 colonies were identified and counted to make a linear curve between CFU/mL and OD.

### Cell isolates and antibiotics

12 clinical MRSP isolates (Cornell Animal Health Diagnostic Center, Bacteriology Laboratory) were grown fresh overnight in CAMHB (BD BBL) broth and diluted down to 2.22*10^6^ CFU/mL using an OD600 reading.

Fresh antibiotic solutions were made in either water or DMSO and diluted in either CAMHB or CAMHB + 2% NaCl. Rifampicin (RIF) (Research Products International) was dissolved in DMSO at a concentration of 20,000 μg/mL and diluted in CAMHB down to 44.4 μg/mL, vancomycin (VAN) (VWR) was dissolved in water at a concentration of 45,000 μg/mL and diluted in CAMHB down to 88.9 μg/mL, linezolid (LIN) (AmBeed) was dissolved in DMSO at a concentration of 60,000 μg/mL and diluted in CAMHB down to 88.9 μg/mL, clindamycin (CLI) (AmBeed) was dissolved in water at a concentration of 80,000 μg/mL and diluted in CAMHB down to 177.8 μg/mL, doxycycline (DOX) (Sigma Aldrich) was dissolved in water at a concentration of 45,000 μg/mL and diluted in CAMHB down to 177.8 μg/mL, and oxacillin (OXA) (Research Products International) was dissolved in water at a concentration of 45,000 μg/mL and diluted in CAMHB + 2% NaCl down to 88.9 μg/mL. Desired concentration ranges were determined by an initial experiment with a separate set of three MRSP strains.

### EZ-SPOTs plates loading and use

On the EZ-SPOTs antibiotic plate, we loaded the plates with a six-slot loader, resulting in five volumes with three replicates for each antibiotic. We also had six regions that were not laser ablated as either a negative or positive control, depending on the corresponding location on the EZ-SPOTs cell plate. Mixing with media results in five two-fold antibiotic concentrations. On the EZ-SPOTs cell plate (Fig. 3B), we loaded one MRSP isolate with a single slot loader and used six regions in the bottom row for negative or positive controls.

### AST experiment

12 96-well microtiter plates were first loaded with 100 μL of CAMHB in columns 1-10 and were loaded with 100 μL of CAMHB + 2% NaCl in columns 11 and 12. Next, antibiotics were loaded on the EZ-SPOTs antibiotic plate and were interfaced with a filled 96-well microtiter plate. After dilution into broth in the 96-well plate, the concentration range for RIF was 0.125 to 2 μg/mL, VAN, LIN, and OXA were from 0.25 to 4 μg/mL, and CLI and DOX were 0.5 to 8 μg/mL. The EZ-SPOTs antibiotics plate was then rinsed with DI water, dried on a paper towel and was reused for each additional isolate. Similarly, the six-slot loader was rinsed with DI water, dried using compressed air and then reused for each additional isolate.

The 96-well microtiter plate containing media and antibiotics was interfaced with the EZ-SPOTs cell plate loaded with one of the MRSP isolates. This results in a 1*10^5^ CFU/mL cell concentration. The EZ-SPOTs cell plate was then soaked in undiluted bleach (Clorox) for two minutes, rinsed with DI water, dried on a paper towel, and reused for each isolate. Similarly, the single-slot loader was soaked in undiluted bleach (Clorox) for two minutes, rinsed with DI water, dried using compressed air, and reused for each isolate.

The resulting 96-well microtiter plates were incubated at 37°C for 24 hours to be consistent with CLSI guidelines. Each plate was then well-mixed and an OD600 reading was obtained for each well using a spectrophotometer to determine cell growth.

### Minimum inhibitory concentration (MIC) determination and clinical breakpoints

After obtaining OD600 growth readings for each 96-well microtiter plate, we averaged replicates, plotted dose-response curves, and determined the minimum inhibitory concentration (MIC), which we defined as the lowest concentration of the antibiotic that completely inhibits bacterial growth in each well. Clinical breakpoints were determined based on CLSI guidelines of susceptibility tests for bacteria isolated from animals.

### Commercially available panel and comparison

These 12 *Staphylococcus pseudintermedius* strains were initially verified by a MALDI Biotyper (Bruker). These strains were run on the Thermo Fisher Sensititre Companion Animal Gram Positive AST plates (COMPGP1F) according to CLSI protocols (Cornell Animal Health Diagnostic Center, Bacteriology Laboratory) and identified as methicillin resistant.

The susceptibility information obtained from COMPGP1F for each of these isolates was kept confidential until the experiment using the EZ-SPOTs plates were completed. The MICs from the Thermo Fisher Sensititre software is the same definition we chose, and thus allowed us to directly compare MICs.

## 4. Conclusion

We demonstrate a robust and easy-to-make low barrier-to-entry solution for performing hundreds to thousands of simultaneous experiments. This platform requires no specialized equipment or knowledge and can be easily made with just a laser cutter. With a few easily obtained materials, this platform is quickly able to be used for a wide range of diagnostic and screening biological and chemical assays. We anticipate that this platform will find broader applications beyond ASTs, particularly in areas that have traditionally lacked high-throughput options.

The platform described in this paper makes it easy to meter and compartmentalize many individual reactions with very high repeatability. One key advantage to the way liquid is loaded with this platform is the decreased chance of a pipetting error in a single or multiple wells. Since all liquid is loaded at once, any human errors would be seen across a whole plate, rather than in a few unexplainable wells. This greatly decreases the amount of time spent following up on false results. Additionally, the plates act as a map or record of the experiment which makes it highly reproducible and readily retraceable.

The ASTs run in this paper would typically require thousands of pipet tips if using a micropipette, but by using EZ-SPOTs we were able to perform the entire experiment with less than one box of pipet tips. The platform can be reused multiple times which further increases the cost-effectiveness, and the lack of pipet tips decreases the biohazardous waste associated with such experiments.

In this paper we interfaced the platform with a 96-well microtiter plate to directly compare our results to the commercially available option. However, by running experiments directly on the EZ-SPOTs plates, the number of individual reactions can easily exceed 1,000 with this footprint or 10,000 with a larger footprint. In these cases, plates can be placed in a humidified incubator or sealed together eliminate evaporation. If interfacing with other microtiter plates is desired, this platform can easily be scaled for well plates containing 384 or 1536 wells.

Such a scalable system allows for an endless possibility of experimental designs, enabling researchers to generate data more quickly. We envision that this versatile platform will be used for many applications including library preparation, drug discovery, protein-based assays, cell-based assays, and solution phase reactions.

## Acknowledgements

We thank the Animal Health Diagnostic Center for initially screening, storing, and providing bacterial isolates. This work was supported by a Cornell Center for Technology Licensing Ignite Innovation Acceleration seed grant and NIH grant P30AI168433. J.A. acknowledges funding support from the Cornell IMSD fellowship and NSF graduate fellowship.

